# Most of Tobacco Male Meiocytes Are Involved in Intercellular Nuclear Migration at the First Meiotic Prophase

**DOI:** 10.1101/2020.07.07.188854

**Authors:** Sergey Mursalimov, Nobuhiko Ohno, Mami Matsumoto, Sergey Bayborodin, Elena Deineko

## Abstract

Serial block-face scanning electron microscopy was used here to study tobacco male meiosis. Three-dimensional ultrastructural analyses revealed that intercellular nuclear migration (INM) occurs in 90–100% of tobacco meiocytes. At the very beginning of meiosis, every meiocyte connected with neighboring cells by more than 100 channels was capable of INM. At leptotene and zygotene, the nucleus in most tobacco meiocytes approached the cell wall and formed nuclear protuberances (NPs) that crossed the cell wall through the channels and got into the cytoplasm of a neighboring cell. The NPs did not separate from the migrating nuclei and never produced micronuclei. Approximately 70% of NPs reached nuclei of neighboring cells. The NPs and the nuclei they reached got very close, and the gap between their nuclear membranes became indistinguishable in some cases. At pachytene, NPs detached from the nuclei of neighboring cells and came back into their own cells. After that, the INM stopped. The reason for such behavior of nuclei is unclear. INM probably causes a short-lived fusion of two nuclei and thus has a potential to form aneuploid or unreduced pollen. We consider INM a normal part of tobacco meiosis.

## INTRODUCTION

Intercellular nuclear migration (INM), also called cytomixis, is an enigmatic phenomenon that can be seen in plant male meiocytes. In this process, nuclei with no signs of damage migrate between cells through special intercellular channels called cytomictic channels (CCs), which are considerably larger than plasmodesmata. INM was discovered more than a century ago and has been so far described in male meiosis of over 400 plant species (for review, see Mursalimov et al., 2013). In most cases, INM can be observed in zygotene and pachytene with highly varied frequency, and it is generally accepted that INM results in the formation of micronuclei (Barton et al., 2014; Reis et al., 2016).

Despite the high prevalence of INM in plant male meiosis, its causes, mechanisms, and consequences are still disputable. On the one hand, INM is regarded by most researchers as a meiotic deviation that does not deserve close attention. Thus, this phenomenon is simply ignored in many works on plant meiosis. On the other hand, some researchers consider INM a process that leads to aneuploid-or unreduced-pollen formation (Farooq et al., 2014; Fakhri et al., 2016; Djafri-Bouallag et al., 2019). This point of view is not supported by sufficient experimental evidence. Attempts to study INM by common cytological techniques have come to a standstill. One of the main obstacles to INM research is inaccessibility of intact male meiocytes for direct analysis because they are hidden inside an anther surrounded by layers of nourishing and protective cells. Common cytological techniques like squashed preparation and tissue sectioning combined with light microscopy (LM) or ultrathin sectioning combined with transmission electron microscopy (TEM) provide only fragmentary information about INM. The mechanical impact on meiocytes on squashed preparations changes their structure and disrupts intercellular channels. In comparison, all kinds of sectioning preserve meiocyte structure but provide only two-dimensional data, and all the information outside a section plane is lost.

Thus, there is a need for new approaches to the study of INM in plant meiosis, and it seems that serial block-face scanning electron microscopy (SBF-SEM) is one of them. SBF-SEM is a relatively new technique that allows for serial sectioning and imaging of resin-embedded material with subsequent digital alignment of hundreds and thousands of ultrastructural pictures and three-dimensional tissue reconstruction. This method was developed to study nervous tissues (Denk and Horstmann, 2004), and there are still just a few examples of this method’s application to plant cell analysis (Kittelmann et al., 2016; Płachno et al., 2017). SBF-SEM has never been employed to study plant meiosis. Nonetheless, it appears that SBF-SEM is the only way to investigate the real picture of such processes as INM that are normally hidden inside organs, can be easily influenced by a mechanical impact, and need to be researched at high resolution.

In this study, we utilized SBF-SEM to investigate INM in tobacco male meiosis. The INM was analyzed in samples where cells were neither mechanically damaged nor lost. For the first time, the exact number of the meiocytes involved in INM was determined, and the number of CCs connecting meiocytes was estimated. We found that 90–100% of meiocytes in a tobacco anther are involved in INM as a nuclear donor, recipient, or both. Furthermore, the migrating nuclei tended to make nucleus–nucleus contacts with the nuclei of neighboring meiocytes. Such nucleus–nucleus contacts seemed to be the only result of INM in tobacco meiosis. No micronuclei were found in the studied cells. The successful observation of unusual behavior of nuclei in tobacco meiocytes—that has been hidden from researchers by the LM and TEM limitations for decades—means that SBF-SEM can open a whole new chapter in plant meiosis research.

## RESULTS

### Most of Tobacco Male Meiocytes Participate in INM

Fourteen anthers, each from an individual tobacco plant, were randomly collected for analysis (Figure 1A). A random tissue fragment from every anther containing meiocytes was analyzed by SBF-SEM. As a result, serial ultrastructural images of tobacco meiocytes were obtained for every sample (Figure 1B). Then, using software, meiocyte nuclei and cell walls were labeled in these images (Figure 1C), and three-dimensional structure of the tissue fragments was reconstructed (Figure 1D and 1E; Supplemental Movie 1).

**Figure 1.**
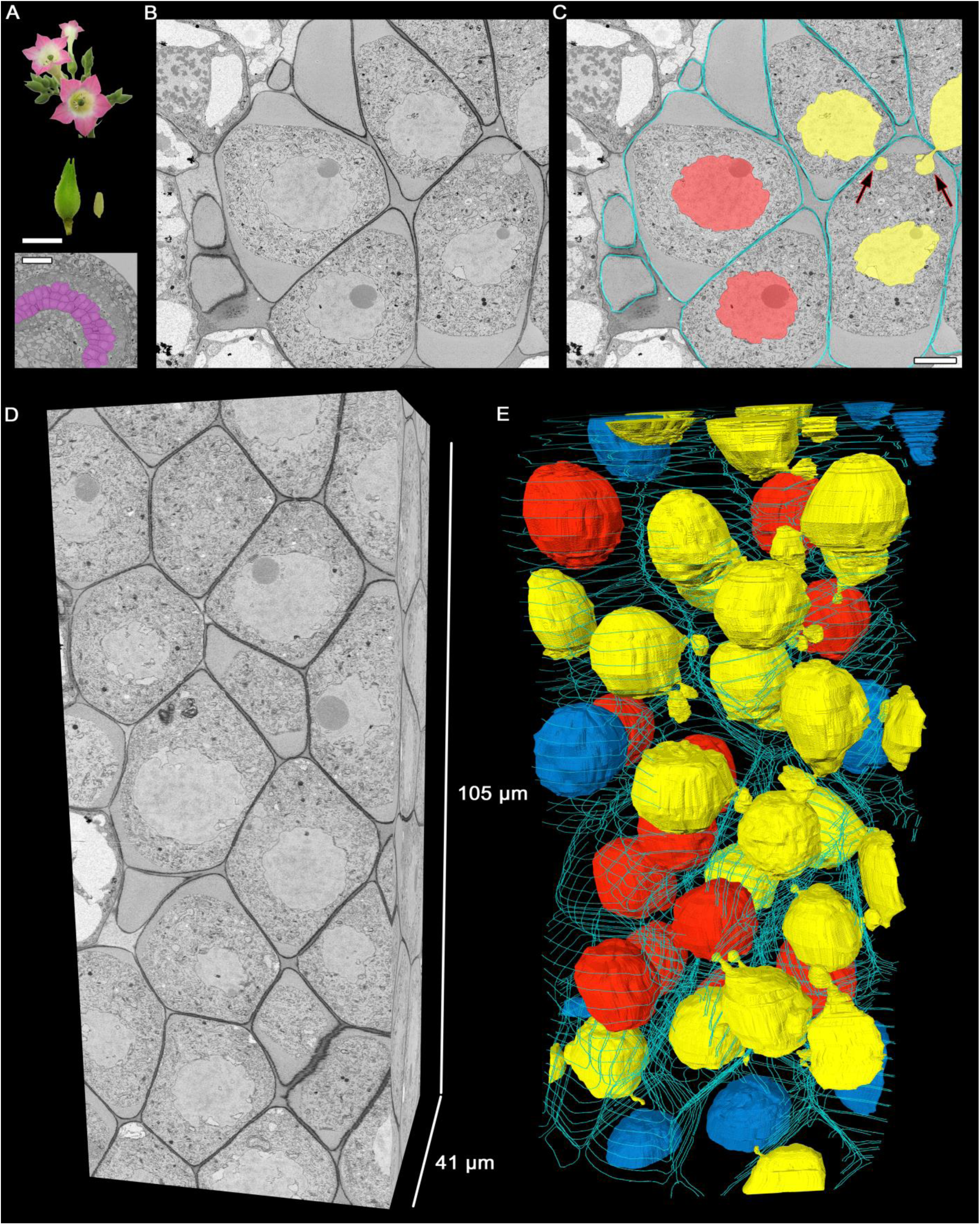
Three-dimensional reconstruction of male meiocytes in a tobacco anther. **(A)** Tobacco flowers, a flower bud, an anther, and transverse section of an anther locule. The purple color indicates meiocytes. **(B)** and **(C)** A meiocyte micrograph obtained by SBF-SEM at pachytene. An original image **(B)** and the same image after cell walls and nuclei were labeled **(C)**. **(D)** and **(E)** Three-dimensional reconstructions of a scanned tissue fragment from 2626 serial images. The whole unlabeled tissue fragment **(D)** and labeled nuclei and cell walls **(E)**. Yellow denotes migrating nuclei, red nonmigrating nuclei, blue partially scanned nuclei (their status is not clear), and turquoise the outer layer of cell walls, labeled on every 50th slice. The arrows point to the NPs crossing a cell wall. Scale bars are 5 mm and 50 µm in **(A)** and 5 µm in **(C)**.

In a typical tobacco meiocyte involved in INM, a nucleus approached the cell wall and formed nuclear protuberances (NPs) that crossed the cell wall through one or a few CCs and got into the cytoplasm of a neighboring cell. Thus, some meiocytes became nuclear donors (Figures 1C and 1E, yellow nuclei). The nuclei of the other meiocytes did not move (Figures 1C and 1E, red nuclei), but these cells could still take part in the INM as recipients of the migrating nuclei. One meiocyte could be both a nuclear donor and recipient at the same time. A nucleus from one donor cell could migrate simultaneously into two recipient cells as well as two nuclei could migrate into one recipient cell (Figures 1C and 1E; Supplemental Movie 1).

We studied tobacco meiocytes at leptotene, zygotene, pachytene, and diplotene of prophase I and at anaphase I (Figure 2A). Special attention was given to zygotene and pachytene. A total of 505 tobacco meiocytes were analyzed, 429 of them were involved in INM and 76 were not (Figure 2B). The observed INM frequency was quite surprising: nearly 100% of meiocytes were involved in the INM as a donor, recipient, or both at certain stages. At leptotene, ∼70% of the tobacco meiocytes participated in the INM. At zygotene, INM frequency increased up to 90–100% in every tobacco anther studied at this stage. At pachytene, it was ∼80–100%, and the INM frequency gradually decreased throughout meiotic stages. At anaphase I, chromatin inside CCs was noted in fewer than 20% of meiocytes. We were unable to find any samples without the INM in the studied material. Besides the high frequency of the INM, we did not find any other visible deviations in the analyzed meiocytes.

**Figure 2.**
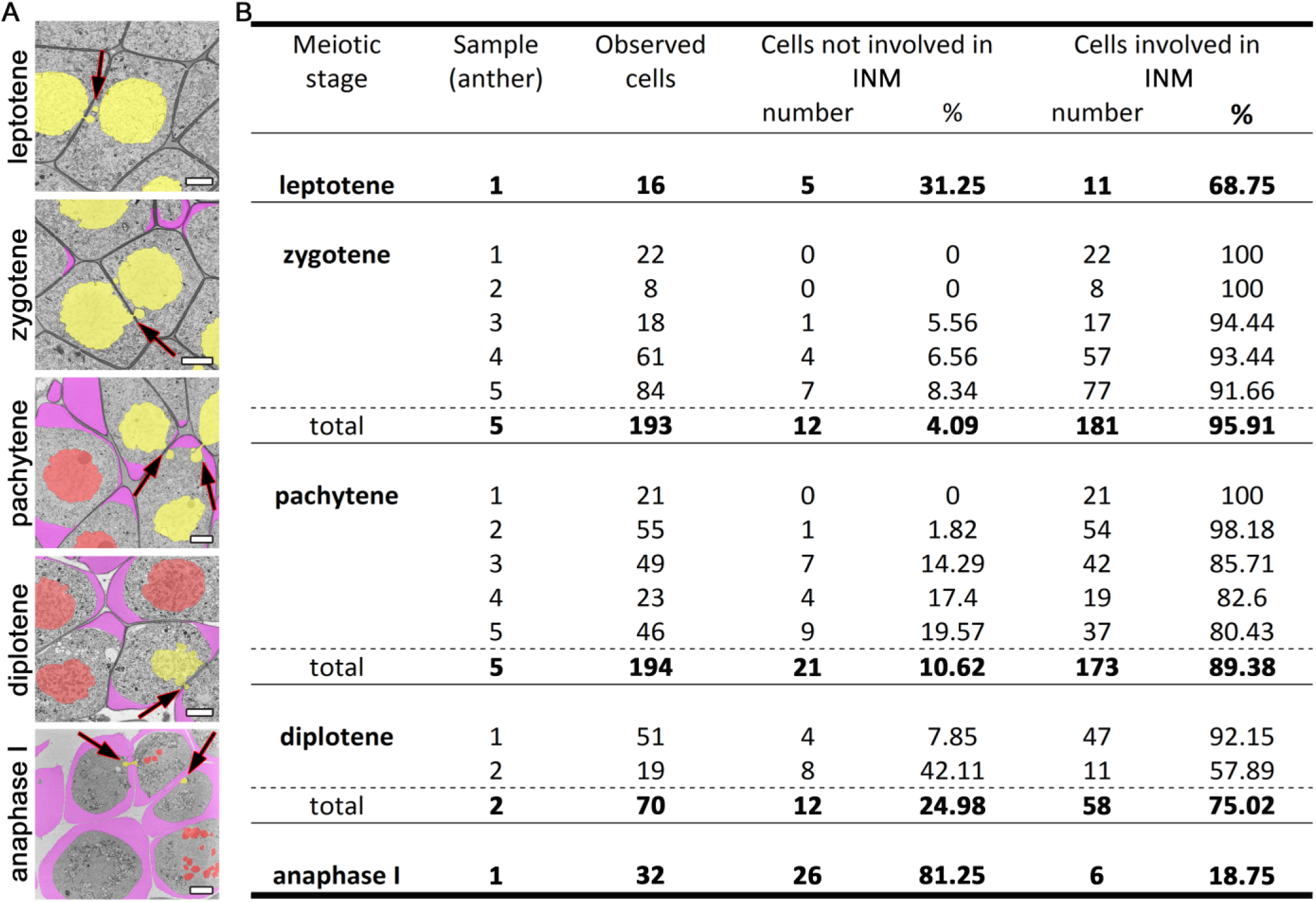
INM frequencies at different meiotic stages. **(A)** Tobacco meiocytes from leptotene to anaphase I. Yellow denotes migrating nuclei, red nonmigrating nuclei, and purple a callose wall; the arrows indicate the NPs crossing a cell wall. **(B)** Statistical data on the INM frequencies. Scale bars are 5 µm.

### The Migrating Nuclei Reach Neighboring-Cell Nuclei

SBF-SEM revealed that in tobacco meiocytes, the migrating nuclei never give rise to micronuclei after the migration. We did not find binucleated meiocytes either. The majority of the observed NPs after crossing the cell wall were headed to the nucleus of the recipient cell still being connected with the nucleus of the donor cell. At zygotene, ∼70% of NPs approached nuclei of recipient cells (Figures 3A to 3C; Supplemental Movie 2). Their nuclear membranes got very close at this stage (Figure 3D, arrowheads), and the gap between membranes of two nuclei became indistinguishable in some cases (Figure 3E, arrowhead). As a rule, NPs formed by the migrating nuclei were oriented in the same direction. Thus, meiocytes often formed chainlike structures connected by NPs (Figures 3A to 3C; Supplemental Movie 2). In most cells, the size of NPs entering the cytoplasm of neighboring cells was ∼3#x2013;5% of the migrating-nucleus volume. Nevertheless, in one meiocyte, we observed two NPs with the size of ∼40% of its nucleus (Figure 4; Supplemental Movie 3). These NPs migrated through two separate CCs but were still located very closely without visible space between their nuclear membranes (Figures 4B and 4C, arrowheads).

**Figure 3.**
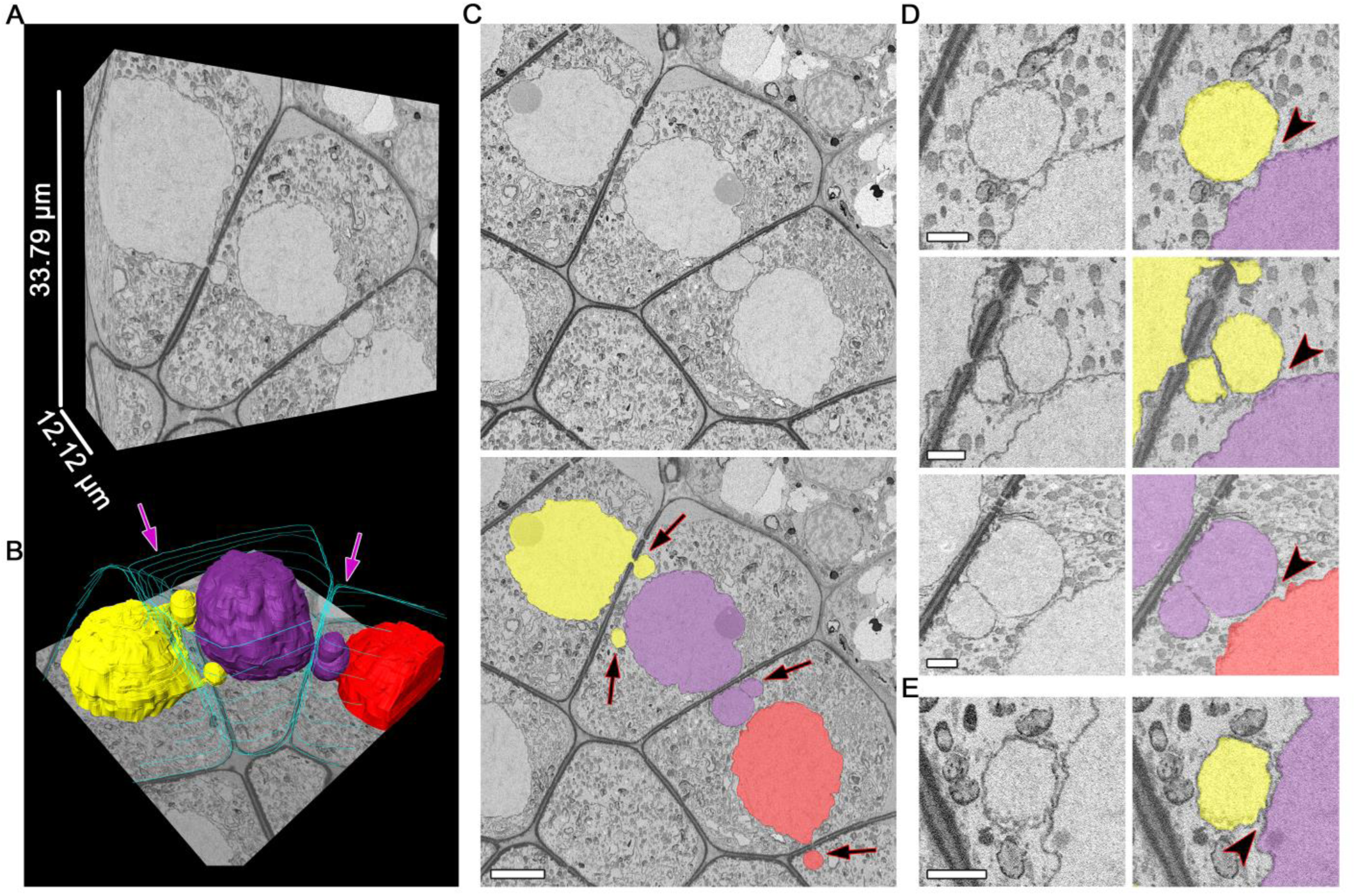
Unidirectional migration of nuclei between meiocytes at zygotene. **(A)** and **(B)** Three-dimensional reconstructions of the scanned tissue fragment by means of 303 serial images. The whole unlabeled tissue fragment **(A)** and labeled nuclei and cell walls **(B)**. **(C)** The original and labeled micrograph. **(D)** and **(E)** Nucleus–nucleus contacts between NPs and nuclei of the neighboring cells, enlarged. The space between nuclear membranes is visible in **(D)** and not visible in **(E)**. Yellow, purple, and red denote individual nuclei (all of them are involved in the INM), and turquoise indicates cell walls, labeled on every 50th slice; the arrows indicate the NPs crossing a cell wall; the arrowheads point to nucleus–nucleus contact areas. Scale bars are 5 µm in **(B)** and 1 µm in **(D)** and **(E)**.

**Figure 4.**
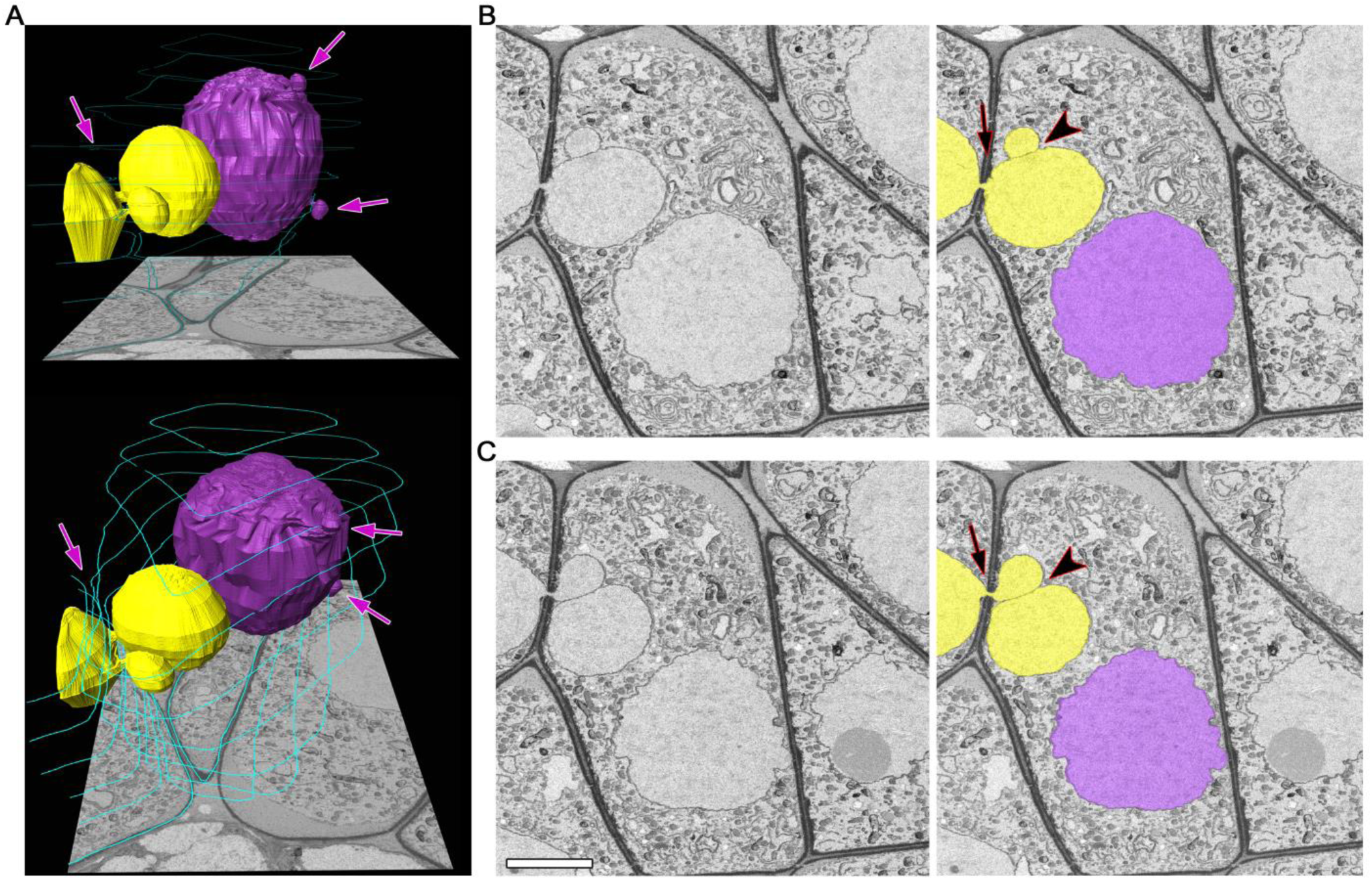
A zygotene meiocyte that has a potential to become binucleated. **(A)** Three-dimensional reconstructions from 464 serial images of labeled nuclei and cell walls in two views at different angles. **(B)** and **(C)** Original and labeled micrographs with the biggest size of the NPs. Yellow and purple denote individual nuclei (both are involved in the INM), and turquoise represents cell walls, labeled on every 50th slice. The arrows indicate the NPs crossing a cell wall, and the arrowheads point to contact areas between the NPs. The scale bar is 5 µm.

### Fewer than 15% of CCs Are Involved in the INM

After three-dimensional reconstruction of individual meiocytes’ structures (Figures 5A to 5C; Supplemental Movie 4), we observed two types of intercellular channels in their cell wall: CCs and plasmodesmata. These types of channels could be identified easily. Plasmodesmata were found to be distributed uniformly in the meiocyte cell wall (Figure 5B). CCs formed clusters in a few cell regions (Figures 5B and 5C). Unlike CCs, plasmodesmata had distinct internal structure (Figure 5E). CCs were more like holes in the cell wall filled with cytoplasm, migrating nuclei, or other organelles such as vesicles or plastids (Figure 5E; Supplemental Movie 5).

**Figure 5.**
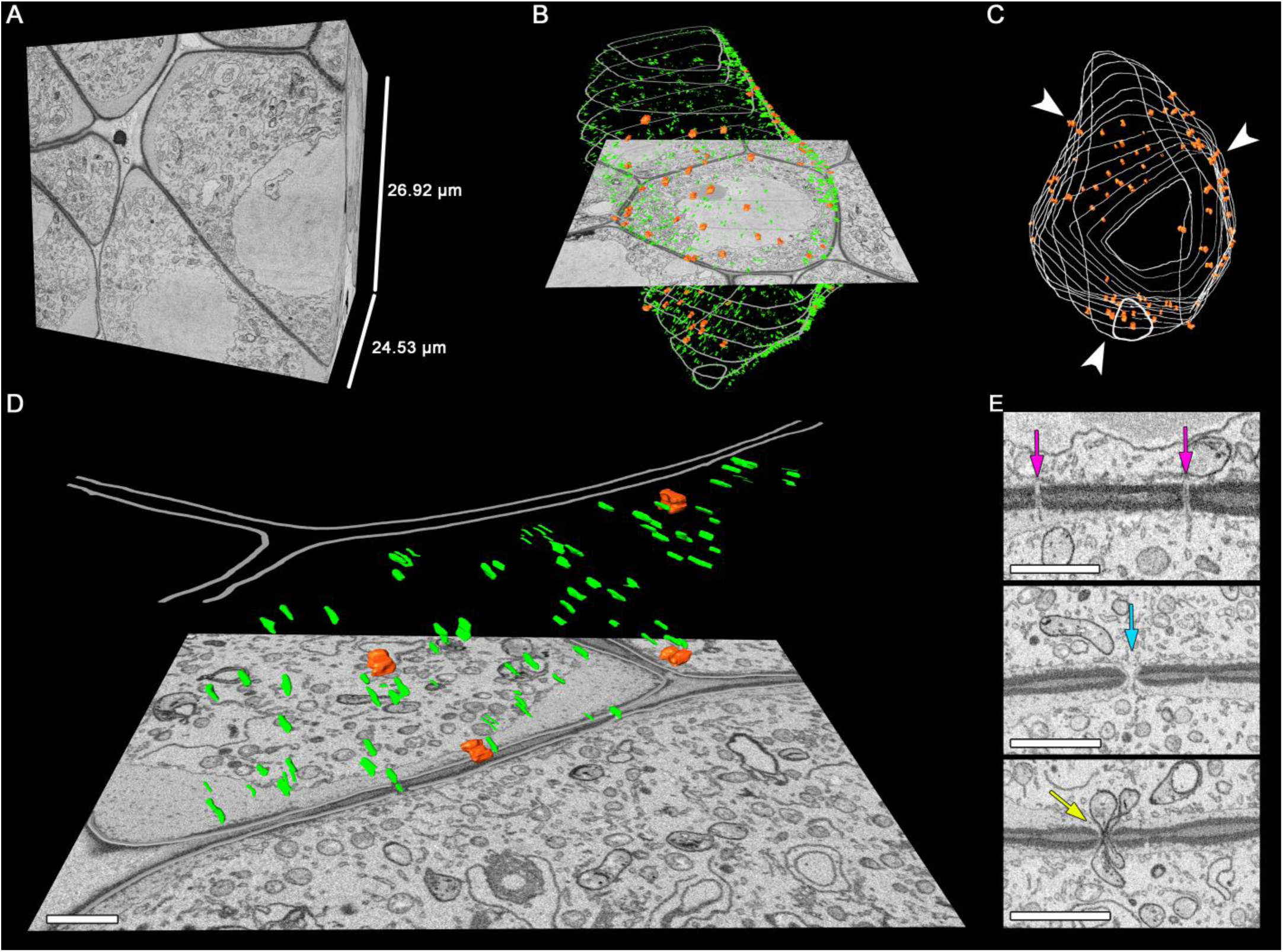
Distribution of the intercellular channels in the tobacco meiocyte cell wall. **(A)** to **(C)** Three-dimensional reconstructions of the scanned tissue fragment from 673 serial images. The whole unlabeled tissue fragment that contains one meiocyte **(A)**, labeled CCs, plasmodesmata, and cell wall **(B)**, with only CCs and the cell wall labeled in the z-projection. **(D)** An enlarged cell wall fragment. **(E)** Sections with plasmodesmata and CCs. Orange represents CCs, green plasmodesmata, and grey the cell wall. Arrowheads indicate the CC clustering regions, the purple arrow points to plasmodesmata on a section, the turquoise arrow indicates a CC on a section filled with cytoplasm, and the yellow arrow points to a CC on a section with two plastids inside. Scale bars are 1 µm.

For the first time, we were able to determine the exact number of CCs that connect tobacco meiocytes (Table 1). At the very beginning of meiosis (leptotene), an average meiocyte had ∼140 CCs in its cell wall. At the following prophase I stages, the CC number decreased quite fast, and at pachytene, an average meiocyte had ∼25 channels. After that, the CC number did not change significantly at least until anaphase I, the last stage we checked. The number of CCs per cell was variable, but their dynamics were obvious in all samples; the highest number of CCs connected tobacco meiocytes at leptotene and then decreased due to callose wall formation (Figure 2A).

**Table 1.**
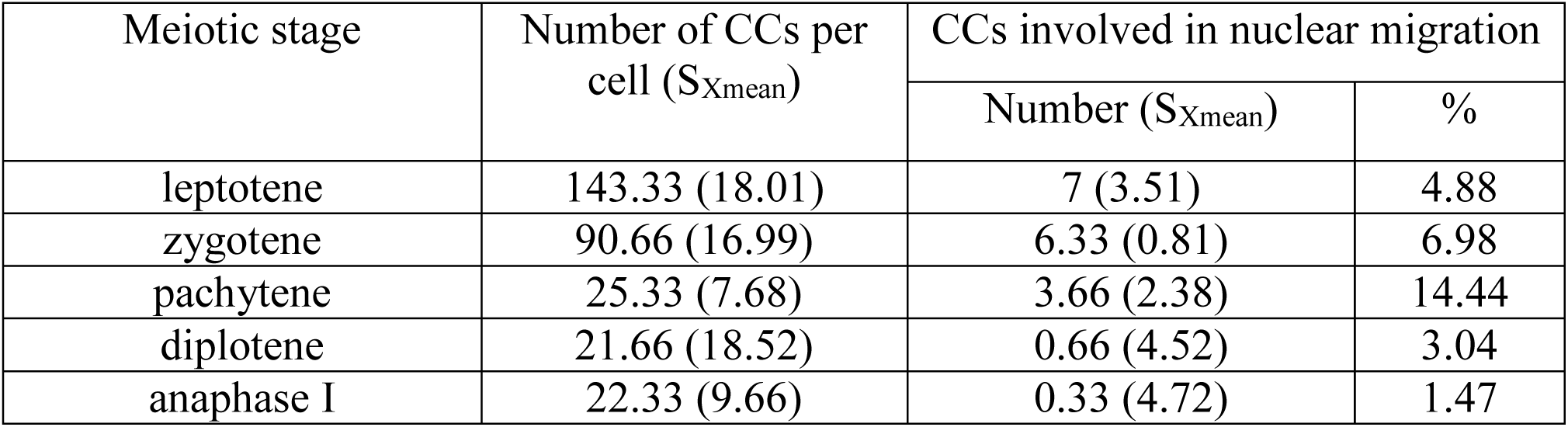
The number of CCs in tobacco meiocytes at different meiotic stages.

Even though tobacco meiocytes contained dozens of CCs, few of them were engaged in the INM. At zygotene and pachytene, when INM frequency was the highest, three to seven CCs had NPs inside in a meiocyte, i.e., fewer than 15% of CCs took part in the INM (Table 1). This number includes both NPs formed by the cell itself and NPs that the cell received from neighboring cells. We observed the formation of CCs only between meiocytes. CCs connecting meiocytes and tapetum cells were not detected.

The plasmodesmata showed dynamics similar to those of CCs during meiosis. Their number was the highest at the leptotene stage and then gradually decreased. Unlike CCs, there were no plasmodesmata left in the meiocyte cell wall at anaphase I.

## DISCUSSION

### INM in Plant Meiosis Is Greatly Underestimated

A common laboratory tobacco line was used in this work. INM is not a unique feature of this line or tobacco in general. INM in male meiosis has been observed in hundreds of angiosperm and in some quillwort species (He et al., 2004; Mursalimov et al., 2013). CCs have been observed in male meiocytes of gymnosperms, ferns, and horsetails (Lehmann et al., 1984; Gabarayeva et al., 2011), and it seems to be only a matter of time for INM per se to be found in these plants. The observed frequency of INM is highly varied among different species and even in one species if it was estimated by different authors. It can vary from less than 1% to 80–100% of male meiocytes in an anther (Li et al., 2009; Reis et al., 2016; Kaur and Singhal, 2019). Such high variation of the numbers makes them doubtful. The only feature that has stayed consistent among many studies is higher INM frequency in the meiosis of polyploid plants (Negron-Ortiz, 2007; Kumar et al., 2011; Reis et al., 2016) and under stressful conditions (Pécrix et al., 2011; Barton et al., 2014). It is important to note that all these estimates have been made by means of squashed preparations combined with LM.

Previously, via the same approach, we studied INM in tobacco plants of different ploidy (Mursalimov et al., 2016). We observed the increase of INM frequency proportionally from 0.6% in diploid plants to 18.6% in triploid and 38.4 % in tetraploid ones. Nevertheless, the INM frequency was estimated at 1.6% in haploid plants, which is not significantly different from that in diploid ones. Diploid plants of the same tobacco line were analyzed in the present study. SBF-SEM revealed that in fact, the INM frequency in their meiosis is ∼90–100%, not 0.6%. Thus, using SBF-SEM, we found that in tobacco meiosis, INM frequency is ∼150-fold higher than estimated before by LM. We believe that there are two reasons for such a big discrepancy. The first reason is the limits of LM resolution: some NPs are too small to be detected this way. The second reason is the mechanical impact on cells on squashed preparations that disrupts NPs and prevents their counting. It is logical to assume that all the studied tobacco plants (haplo-, di-, tri-, and tetraploids) have the same INM frequency, and the only reason we see differences in estimated numbers is the use of an inappropriate method for this analysis (squashed preparations combined with LM). It seems that the amount of migrating chromatin is critical for INM detection by LM. In other words, the more chromatin migrates between cells, the easier it is to detect. This notion explains the observed higher INM frequency in tri-and tetraploid tobacco plants. The amount of the migrating chromatin is greater at every subsequent ploidy level. The larger chromatin amount binds more dye molecules. Therefore, the migrating chromatin becomes more visible and can be registered readily. In this context, it is not surprising that the INM frequency in haploid plants is not significantly higher because the chromatin amount is not greater either, and probably only the biggest NPs can be detected by LM in this case. These conclusions may be valid for polyploid forms of other plant species too.

The enhancement of chromatin stainability can also be the reason for the observed high INM frequency in the meiosis of plants under stressful conditions. It is known that exposure to stressors such as abnormal temperature or radiation can cause the death of meiocytes and sterility (De Storme and Geelen, 2014). Chromatin is more condensed and stains more intensely with dyes in dying plant cells than in live cells (Yamada et al., 2006). The migrating chromatin in the dying meiocytes is also more condensed and more strongly stained; therefore, it may be detected more easily by LM. This is a possible reason for the high INM frequency observed in plants under stressful conditions.

Accordingly, we assume that INM frequency is high in normal meiosis of many plant species and is not influenced by ploidy or stress. Nonetheless, it may be fully detected by LM only in polyploids that have a large amount of migrating chromatin or after stress exposure in dying meiocytes when the migrating chromatin gets condensed and overstained.

It is still too early to state that the majority of male meiocytes are involved in INM in all plant species where it has been detected. On the other hand, it is obvious that INM has been underestimated so far, and in light of our results, all the published data about INM frequency have to be reconsidered.

### INM Never Gives Rise to Micronuclei in Tobacco Meiocytes

It is generally accepted that micronucleus formation is the main consequence of INM in all the plant species where INM has been detected including tobacco (Mursalimov and Deineko, 2011; Barton et al., 2014; Reis et al., 2016). Here, SBF-SEM revealed that this point of view is incorrect. INM never produced micronuclei in tobacco meiosis. It is possible that in the meiosis of other plant species, INM does not produce micronuclei either. Previous incorrect conclusions about the INM consequences in tobacco and possibly in other species have been made due to misinterpretation of the results of LM and TEM of squashed preparations and sections. As mentioned above, these techniques’ limitations lead to the underestimation of INM frequency, and apparently for the same reasons, artificial micronuclei are observed. Many of NPs inside CCs can be disrupted on squashed preparations. Even if some of them persist, they are often too small to be detected by LM. When INM is studied by TEM, NPs inside CCs may be invisible outside the plane of an ultrathin section. By contrast, NPs that already left CCs and entered the cytoplasm of a recipient cell are more prominent due to their bigger size. This set of NPs cannot hide from analysis so easily. This state of affairs leads to the observation of the artificial micronucleuslike structures. In our study, SBF-SEM revealed that NPs inside CCs are not disrupted in the absence of a mechanical impact, and NPs located in the cytoplasm of recipient cells do not lose the connection with the main part of their nucleus staying in a donor cell.

There are three other possible scenarios for the migrating nuclei if they do not produce micronuclei: (i) a whole nucleus can cross the cell wall without dividing into parts and form a binucleated cell, (ii) NPs can fuse with a nucleus of a recipient cell and become undetectable, and (iii) NPs can return into the donor cell without any consequences. The formation of binucleated meiocytes as a result of INM has been described in several plant species. It has been shown that a whole nucleus can migrate from a donor to recipient cell, thereby forming one binucleated and one empty meiocyte. Presumably, such binucleated meiocytes can produce unreduced pollen (Singhal and Kumar, 2008; Tsvetova and Elkonin, 2013; Sidorchuk et al., 2016). The frequency of binucleated-meiocyte formation as a consequence of INM has not been estimated before for any plant species. In our study, SBF-SEM revealed that in tobacco meiosis, one in 429 meiocytes involved in INM has a potential to become a binucleated cell. Thus, due to its low frequency, the binucleated-meiocyte formation cannot be the main consequence of the INM in tobacco.

The observed by SBF-SEM tendency for the establishment of nucleus–nucleus contacts during the INM suggests that nuclear fusion is a more likely consequence of this process. Nuclear fusions during INM in tobacco male meiocytes have already been documented by TEM. After the fusion, two nuclei stay in their own cells but make contact through CCs and share one nuclear membrane forming a bridgelike structure filled with chromatin (Mursalimov and Deineko, 2011). Nonetheless, only a few cases of such bridgelike-structure formation have been registered by TEM, and they have never been detected by LM in tobacco or by LM/TEM in other species. Here, SBF-SEM indicated that most nuclei in tobacco meiocytes are prone to fusion. Although we did not directly observe nuclear fusions by SBF-SEM, we noted numerous cases of the nucleus–nucleus contacts that can transform into nuclear fusions. It can be hypothesized that distinct bridgelike structures between nuclei exist for a very short period, and that is why it is difficult to detect them. To prove that the observed nucleus–nucleus contacts transform into nuclear fusions, INM should be studied in live meiocytes at resolution comparable with that of electron microscopy. Unfortunately, there are no suitable methods right now, but new approaches to plant meiosis research are being developed quite fast, and we expect to have new tools to study the INM in plant meiosis in the nearest future.

Taken together, our results suggest the following preliminary interpretation of the observed phenomenon. The main purpose of INM is the establishment of nucleus–nucleus contacts. To this end, nuclei migrate through CCs at leptotene and zygotene and are headed to neighboring nuclei. Then, they fuse forming short-lived bridgelike structures. At pachytene, the nuclei lose their connection, the migrating nuclei return to their initial position in donor cells, and the meiocytes continue meiosis independently. It appears that meiocytes complete the INM before the nuclear-membrane disappearance at the end of the first meiotic prophase. The presence of chromatin inside CCs in later meiotic phases such as anaphase I is most likely a deviation from typical INM. In this case, migrating nuclei probably did not manage to return to the donor cells completely before their nuclear membrane disappeared, and the chromatin got stuck inside CCs.

It is possible that nucleus–nucleus interactions during INM result in changes of the chromatin amount in nuclei of both donor and recipient cells. The nuclei can gain or lose some chromatin/chromosomes. As a result, pollen with altered karyotype will be formed. It is well known that polyploidization is one of the major driving forces behind plant evolution, and the unreduced- or aneuploid-pollen formation plays a key role in this process (De Storme and Mason, 2014). In contrast to animals and yeast, plant meiosis is a rather robust process that has no critical checkpoints (Wijnker and Schnittger, 2013). Accordingly, plant meiotic division continues even if the chromosome number changes or their homologous pairing is absent. In this context, INM can be an additional way to increase genetic diversity because of the formation of aneuploid or unreduced pollen. It is noteworthy that INM is seen in plant meiocytes at the same meiotic stages when crossing over occurs. This means that meiocytes are primed for genetic recombination at this moment.

Obviously, our interpretation of the observed phenomenon requires additional experimental evidence probably obtained by approaches that have not been developed yet. We have no other explanation for the observed picture of events. It can be theorized that the observed INM and nucleus–nucleus contacts have no consequences, and the migrating nuclei simply return to their cells unchanged. But for what reason do they do so in this case? Why do these cells put so much effort into a useless process? On the other hand, it is impossible to explain all the obtained data by some kind of meiotic aberrations or some artificial influences.

In this study, SBF-SEM also revealed that CC functions are not limited to INM. In fact, only a few CCs are involved in this process in every meiocyte. In addition, there are some meiocytes in an anther that contain dozens of CCs and do not participate in INM at all. Due to the possibility of free movement of cytoplasm and small organelles through CCs, their main function may be the transfer signaling molecules and nutrients that is essential for synchronous meiotic division.

Thus, SBF-SEM indicates that the INM in tobacco male meiosis has been greatly underestimated so far. INM, which has been regarded as a meiotic deviation by many authors, in reality, seems to be a constant part of normal tobacco male meiosis. In other words, the absence of INM in tobacco meiocytes is likely to be a deviation. Considering the high prevalence of INM in meiosis among various plant species and its similar patterns in all observed cases, we believe that the INM picture observed in tobacco is relevant for all the plant species where INM has been detected. This means that in many/all higher plant species, the majority of their male meiocytes can take part in INM, leading to short-lived nuclear fusions. Of course, this is only a hypothesis right now. Nonetheless, it is amazing how the real picture of INM in plant meiosis has been hiding in plain sight for decades owing to the limitations of LM and TEM. Definitely, more researchers must try to tackle this problem, thereby starting a whole new chapter in the study of plant meiosis by SBF-SEM and other new methods.

## METHODS

### Plant Growth and Sampling

A wild-type tobacco line (*Nicotiana tabacum* L. cv. Petit Havana SR1) with high pollen fertility was used in this work. The plants were grown in a greenhouse with a photoperiod of 16/8 h (day/night) at a temperature of 22/18°C (day/night). One anther per plant (14 total) was randomly chosen for analysis.

### Observations and Image Analyses by SBF-SEM

Anthers were fixed with 2.5% paraformaldehyde and 2.5% glutaraldehyde in 0.08 M cacodylate buffer (pH 7.2) on ice overnight. After that, the tissue was washed three times for 15 min with 1× phosphate-buffered saline (PBS, pH 7.2). Next, the anthers were mounted in 3% low-melting-point agarose in 1× PBS and cut on a microtome with a vibrating blade (HM 650V; Microm, Germany) into 200 µm pieces. Then, they were postfixed with a mixture of 1.5% potassium ferrocyanide and 2% osmium tetroxide in 1× PBS for 1 h at 4 °C. The samples were next washed four times for 3 min in MilliQ water at room temperature and incubated in a 1% solution of thiocarbohydrazide (Sigma-Aldrich, USA) for 20 min at 60 °C. After that, the samples were washed four times for 3 min in MilliQ water and placed in 2% aqueous osmium tetroxide for 30 min incubation, then the tissue was washed again four times for 3 min in MilliQ water and incubated overnight in 2% aqueous uranyl acetate at 4 °C. The samples were then rinsed four times for 3 min in MilliQ water, incubated in a freshly prepared Walton’s lead aspartate for 2 h at 50 °C, washed four times for 3 min in MilliQ water, and dehydrated in 30%, 50%, 70%, and 96% ethanol solutions for 10 min each, one time in 4°C 96% ethanol for 10 min, and then three times for 20 min in acetone at room temperature. After the dehydration, the samples were placed in a mixture of Araldite (Fluka, Switzerland) epoxy resin and acetone in ratios 1:2, 1:1, and 2:1 in that order, with incubation for 1 h at each step at room temperature. Then, the samples were kept in 100% resin overnight at room temperature and polymerized over 2 nights at 60 °C.

The polymerized samples were cut out and glued with silver-containing conductive paste (KAKEN TECH, Japan) onto aluminum specimen pins and evaporated with gold in vacuum. Serial images were acquired with a ΣIGMA field emission scanning electron microscope (Carl Zeiss, Germany) equipped with 3View, a system with a built-in ultramicrotome and a back-scattered electron detector (Gatan, USA), at a resolution of 6.3 nm/pixel in X and Y directions and 40 nm in the Z direction. The size of the imaged area was 51.6096 × 51.6096 µm, which resulted in an image resolution of 8192 × 8192 pixels. From 387 to 3689 serial images were obtained from every sample. The images from each dataset were processed in the Fiji software (Schindelin et al., 2012). Manual image segmentation was performed with Microscopy Image Browser (Belevich et al., 2016). Three-dimensional tissue reconstruction and its analyses were performed in Amira 6.2.0 software (FEI Visualization Science Group, Hillsboro, OR, USA). For CC number estimation, three meiocytes were chosen randomly for every studied meiotic stage.

## ACKNOWLEDGMENTS

We thank Atsuko Imai, Nobuko Hattori (National Institute for Physiological Sciences), and Tatyana Shnayder (Institute of Cytology and Genetics) for their technical assistance. The material preparation at the Joint Access Center for Microscopy of Biological Objects was supported by ICG project No. 0324-2019-0042-C-01. The use of the Electron Microscopy Unit at the National Institute for Physiological Sciences was supported by Cooperative Study Programs of the National Institute for Physiological Sciences. The English language was corrected and certified by shevchuk-editing.com.

## AUTHOR CONTRIBUTIONS

S.M. designed and coordinated the study, prepared the material, interpreted the data, drafted the manuscript; N.O. participated in interpretation of the data and drafting the manuscript; M.M. performed microscopy analysis; S.B. participated in the material preparation, interpreted the data and E.D. initiated the study, interpreted the data, revised the manuscript critically.

---

**Supplemental Movie 1**. Three-dimensional reconstruction of male meiocytes at pachytene in a tobacco anther.

Reconstructed from 2626 serial images. Yellow denotes migrating nuclei, red nonmigrating nuclei, blue partially scanned nuclei (their status is not clear), and turquoise the outer layer of cell walls, labeled on every 50th slice.

**Supplemental Movie 2**. Unidirectional migration of nuclei between meiocytes at zygotene. Reconstructed from 303 serial images.

Yellow, purple, and red denote individual nuclei (all of them are involved in the INM), and turquoise indicates cell walls, labeled on every 50th slice.

**Supplemental Movie 3**. A zygotene meiocyte that has a potential to become binucleated. Reconstructed from 464 serial images. Yellow and purple denote individual nuclei (both are involved in the INM), and turquoise represents cell walls, labeled on every 50th slice.

**Supplemental Movie 4**. Distribution of the intercellular channels in the tobacco meiocyte cell wall.

Reconstructed from 673 serial images. Orange represents CCs, green plasmodesmata, and grey the cell wall.

**Supplemental Movie 5**. An enlarged cell wall fragment.

Reconstructed from 130 serial images. Orange represents CCs, green plasmodesmata, and grey the cell wall.

